# Identifying Genetic Regulatory Variants that Affect Transcription Factor Activity

**DOI:** 10.1101/2022.10.21.513166

**Authors:** Xiaoting Li, Tuuli Lappalainen, Harmen J. Bussemaker

## Abstract

Assessing the functional impact of genetic variants across the human genome is essential for understanding the molecular mechanisms underlying complex traits and disease risk. Genetic variation that causes changes in gene expression can analyzed through parallel genotyping and functional genomics assays across sets of individuals. Trans-acting variants are of particular interest, but more challenging to identify than cis-acting variants. Here, to map variants that impact the expression of many genes simultaneously through a shared transcription factor (TF), we use an approach in which the protein-level regulatory activity of the TF is inferred from genome-wide expression data and then genetically mapped as a quantitative trait. To analyze RNA-seq profiles from the Genotype Tissue Expression (GTEx) project, we developed a generalized linear model (GLM) to estimate TF activity levels in an individual-specific manner. A key feature is that we fit a beta-binomial GLM at the level of pairs of neighboring genes in order to control for variation in local chromatin structure along the genome and other confounding effects. As a predictor in our model we use differential gene expression signatures from TF perturbation experiments. We estimated genotype-specific activities for 55 TFs across 49 tissues and performed genome-wide association analysis on the virtual TF activity trait. This revealed hundreds of TF activity quantitative trait loci, or aQTLs. Altogether, the set of tools we introduce here highlights the potential of genetic association studies for cellular endophenotypes based on a network-based multi-omic approach.

## INTRODUCTION

In recent years, there has been a large effort to understand the phenotypic impact of genetic variation via genome-wide association studies (Claussnitzer et al., 2020) (Welter et al., 2014). The majority of variants detected by GWAS are noncoding, which makes it difficult to uncover the underlying molecular mechanisms (Ward & Kellis, 2012) (Maurano et al., 2012). Genetic variation often modulates cellular phenotypes through changes in gene expression (Cookson et al., 2009) (Nicolae et al., 2010) (Pickrell et al., 2010). Genetic variants that influence the expression level of a gene are known as expression quantitative trait loci (eQTLs) (Brem et al., 2002). They can impact gene expression either via cis-acting (proximal) or trans-acting (distal) mechanisms (Grundberg et al., 2012) (THE GTEX CONSORTIUM, 2020) (Morley et al., 2004) (Westra et al., 2013) (Võsa et al., 2021).

To date, most studies have focused on identifying cis-acting regulatory variants using eQTL analysis. Mapping of trans-acting genetic variants to specific downstream genes is limited by statistical power because genome-wide tests come with a burden of multiple testing, and trans-eQTLs tend to have a smaller effect size (Yvert et al., 2003) (THE GTEX CONSORTIUM, 2020). Trans-acting variants in principle can affect a large number of genes by altering the activity of gene regulatory pathways (Brynedal et al., 2017) (Hansen et al., 2008). Indeed, mapping the genetic determinants of (inferred) transcription factor (TF) activity as so-called activity quantitative trait loci (aQTLs) was previously shown to be a viable strategy for mapping trans-acting loci in model organisms (Lee & Bussemaker, 2010). In the regulatory network of the cell, the aQTL can be connected to the TF through a variety of mechanistic paths, although it by definition is upstream of the TF in the network in a causal sense. For instance, a causal gene near the aQTL could encode a co-factor of the trait TF, a kinase that controls the post-translational modification status of the TF, or even an enzyme whose modulation leads to a change in metabolic state that gets sensed by a signaling pathway upstream of the TF.

The recent emergence of large collections of parallel genotype and RNA-seq expression data (THE GTEX CONSORTIUM, 2020) has put human aQTL analysis using a similar discovery approach within reach, and some initial studies have been performed (Paull et al., 2021) (J. C. Chen et al., 2014) (Hoskins et al., 2021). Identifying aQTLs could be important both for understanding how genetic variants affect not only molecular phenotypes in *cis* or complex diseases in GWAS, but also intermediate phenotypes of cellular regulatory systems. The protein-level regulatory activity of a TF quantifies to what extent the TF can impact the expression of target genes in a given cell. Most of the current experimental methods for protein quantification only measure the total protein abundance for a TF (Uhlén et al., 2015). However, the activity of a TF protein is greatly influenced by its post-translational modification status and consequent subcellular distribution. Linear regression models have been used to estimate protein-level TF activity from genome-wide mRNA expression levels in a sample-specific manner (Gao et al., 2004) (Bussemaker et al., 2001) (Foat et al., 2006). In such analysis, observed mRNA expression levels serve as response variable and the regulatory influence of a given TF on each gene is used as predictor; the regression coefficient, therefore, reflects the (inferred) activity of the TF under a certain condition. A common way to define a TF’s regulatory influence is through the prediction of binding affinity from promoter sequence using TF binding motifs (Bussemaker et al., 2001) (Conlon et al., 2003) (Schaid et al., 2018) (Balwierz et al., 2014) (Li et al., 2014), or a binary approach based on so-called TF regulons (Schubert et al., 2018). Other computational methods have also been proposed to infer TF activity from the mRNA expression levels of TF target genes (Schaid et al., 2018) (Balwierz et al., 2014) (Li et al., 2014) (Barenco et al., 2009) (Y. Chen et al., 2017) (Fröhlich, 2015) (Fu et al., 2011) (Jiang et al., 2015) (Khanin et al., 2007) (Nachman et al., 2004) (Sanguinetti et al., 2006) (Schacht et al., 2014) (Conlon et al., 2003) (Boulesteix & Strimmer, 2005).

One of our goals in this study was to update the linear regression framework to allow for optimal analysis of human RNA-seq data such as that generated by the GTEx project. First, because it is much more difficult to predict cis-regulatory logic from non-coding sequence in human than in model organisms, we used the observed response of each gene after perturbation of a TF in a cell line as the predictor variable. This reflects the effect of TF regulation on gene expression more accurately than binding affinity predicted from sequence or in terms of TF regulon (the set of target genes) membership. A second issue we needed to address was that especially in human cells, local chromatin context shows great variation along the genome, and has a large influence on gene expression levels (Trescher et al., 2017). While some algorithms (Li et al., 2014) (Jiang et al., 2015) include multi-omics data as input to remove the confounding effect of chromatin context, matched datasets would not be readily available when analyzing genetic variation in gene expression. The solution we settled on is to analyze gene expression data at the level of *pairs* of neighboring genes, which are more likely to be embedded in the same local chromatin type. More generally, our gene-pair approach allows us to control for confounders that affect inter-individual variation in gene expression in complex tissue data. To make optimal use of the discrete RNA-seq counts as a dependent variable, and properly account for the over-dispersion that the distribution of these counts is known to be exhibit (Robinson et al., 2010), we fit a generalized linear model (GLM) based on the beta-binomial distribution with an independent over-dispersion parameter for each gene pair. By combining all these ingredients, our approach provides an insight into how TFs regulate differential gene expression across tissues and individuals. To understand how genetic variation affects gene expression via trans-acting mechanisms by impacting related TF activities, we then performed genome-wide association analysis with the inferred TF activity as a quantitative trait to identify genetic variants (aQTLs) that are significantly associated with TF activity levels in each tissue.

## RESULTS

### Inferring individual-specific transcription factor activity

**Figure 1a** shows an overview of our approach to mapping trans-acting loci. One of the inputs are gene expression data in the form of raw RNA-seq counts for each sample as profiled in the GTEx project v8 release (THE GTEX CONSORTIUM, 2020). These data play the role of dependent variable (response) in the generalized linear model (GLM) that we fit to infer sample-specific, protein-level TF activities (see **Supplemental Figure S1a** and **Methods** for details). The other input is a genome-wide signature, consisting of the log-fold change in expression for all genes after CRISPRi knockdown of a particular TF (**Figure 1b**) derived from RNA-seq data from the ENCODE portal (Dunham et al., 2012), which serves as the independent variable (predictor) in the model. Of the 74 TFs for which such experiments were performed, 55 passed our quality control criterion (see **Methods**).

**Figure 1:**
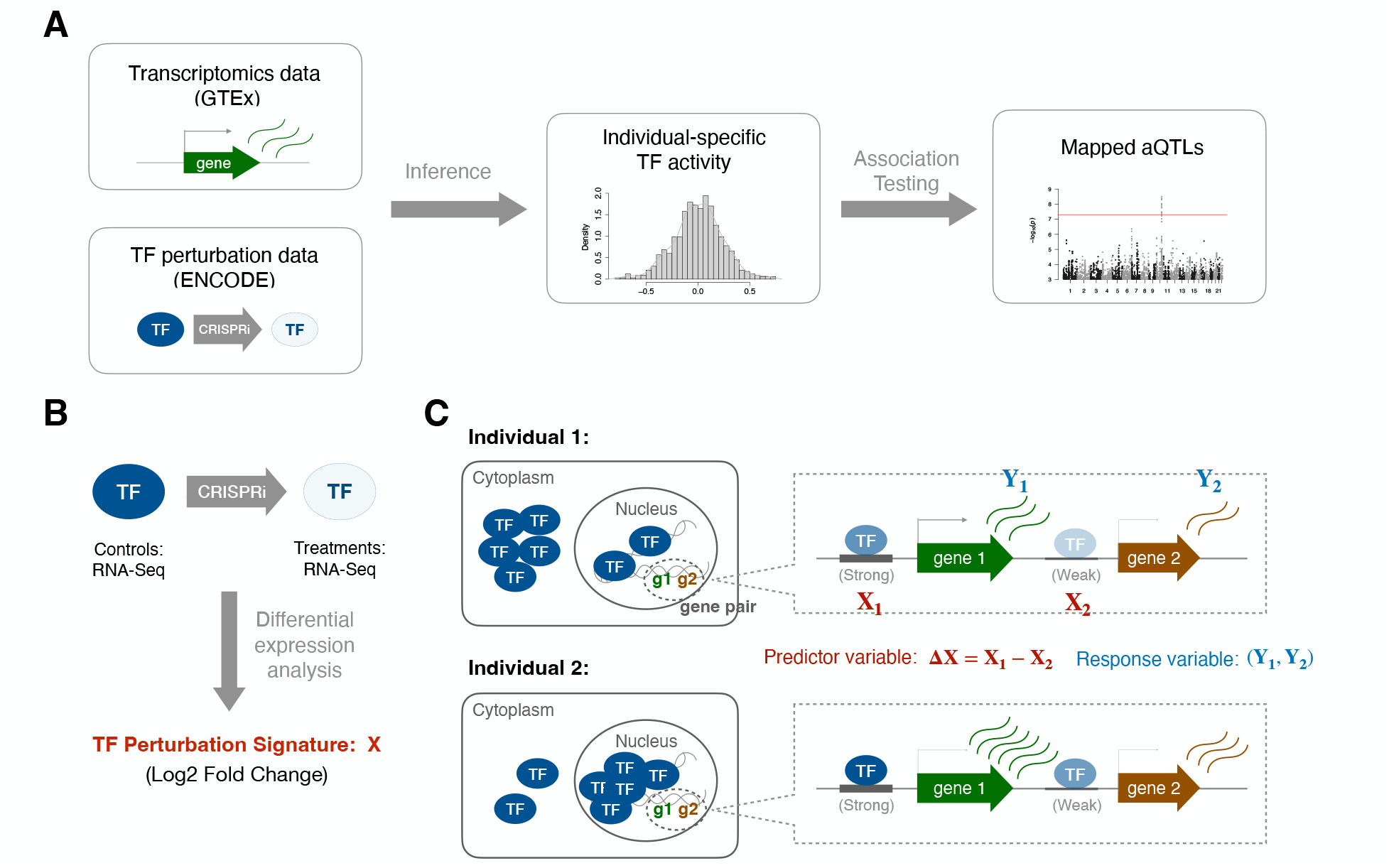
Inferring individual-specific TF activity levels. (a) Overview of the aQTL mapping pipeline. (b) TF perturbation response signature is used as the TF regulatory strength. (c) Diagram of the pair-level model.

In existing regression models used for inferring TF activity from expression data (Gao et al., 2004) (Schaid et al., 2018) (Balwierz et al., 2014) (Li et al., 2014) (Jiang et al., 2015), each observation is an absolute or differential expression value for an individual gene. The regression coefficient of the model in this setup would quantify to what extent the sensitivity of genes to CRISPRi perturbation of the TF is predictive of differential mRNA expression in an unrelated sample. To account for the confounding effect of variation in local chromatin context along the genome without the need to explicitly add covariates related to chromatin context to our model, we developed a “pair-level” model in which each observation is a pair of expression values for two neighboring genes (**Figure 1c** and **Methods**). The rationale is that since chromatin is organized into domains along the genome (Dixon et al., 2016), neighboring genes are more likely to share a similar chromatin environment. Our model is only trying to explain how sample-specific variation in the expression ratio between the two genes in the pair can be explained in terms of differences in responsiveness to TF knockdown between the two genes in the pair. This model definition implicitly accounts for, and therefore is insensitive to, any increase or decrease in expression resulting from local chromatin context that would be shared among the two genes in the pair, Including technical and biological nuisance variation that is inherent to tissue RNA-seq data. Pairs of neighboring protein-coding genes were selected for inclusion in the model based on the distance between their respective transcription start sites (**Supplemental Figure S2a**). We imposed both a minimum distance (10 kb) to avoid sharing of promoter or proximal enhancer regions, and a maximum distance (100 kb) to increase the odds of similarity in local chromatin context.

To properly account for the biological and technological components of the variation in RNA-seq count, we use a GLM based on the beta-binomial distribution and likelihood maximization across hundreds of samples for a given tissue to estimate the dispersion parameter for each observation directly from the data. Simultaneous estimation of the regression coefficient associate with the TF perturbation signature yields an estimate of (differential) TF activity for each sample (see **Methods** for details).

To assess the robustness of our approach, we compared the consistency in inferred TF activity between fits based on odd pairs and even pairs of genes (**Supplemental Figure S2b**). For a representative set of TFs, the Pearson correlation across samples between TF activities based on odd pairs and even pairs was greater than 0.9 (**Supplemental Figure S2c**,**d**). This analysis also provided us with an opportunity to assess to what extent the pairwise nature of our model affected the estimation of TF activities. Consistent with our expectation, when we performed fits in which the pairs are non-nearest neighbors at a larger distance from each other on average, the inferred TF activities were less similar to those inferred based on nearest-neighbors pairs, and more similar to those inferred using individual-gene observations (**Supplemental Figure S3**).

To assess to what extent the variation in inferred TF activities reflected true differences between samples as opposed to statistical fluctuations or biological noise unrelated to the TF in question, we randomly permuted the TF perturbation response signature used as the predictor and refit the model (**Supplemental Figure S1b** and **Methods**). Comparing the null distribution of TF activities resulting from many independent such permutations with the distribution across samples for the unpermuted fits, we found that the variance of the latter tended to be much larger (a representative example is shown in **Figure 2a**). Indeed, a variance ratio between (i) the difference between observed variance and permuted variance and (ii) the observed variance is greater than 0.9 for most TF/tissue combinations (**Figure 2b**). This suggests that our modeling approach is sensitive to true variation in TF activity across samples, which can arise from many sources, including inter-individual variation due to donor characteristics.

**Figure 2:**
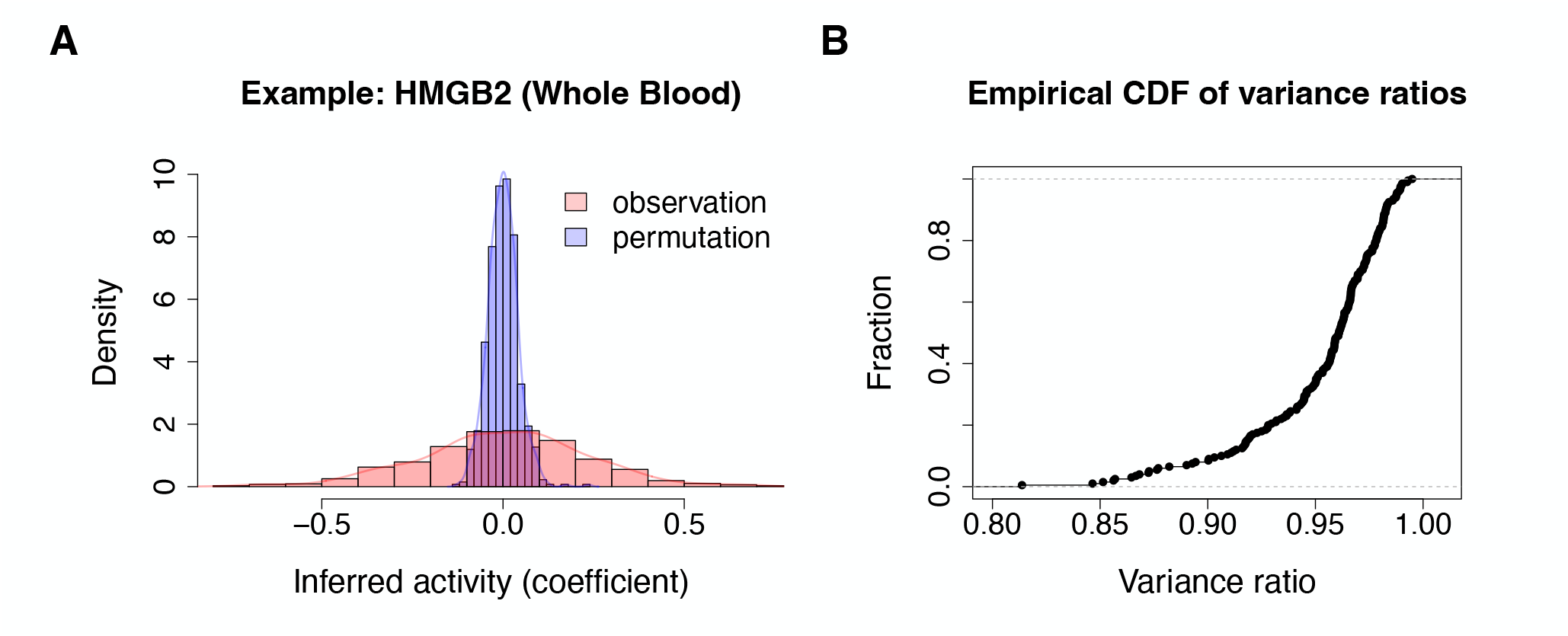
Permutation analysis. (a) An example of distributions of observed inferred activity and permuted inferred activity. (b) Distribution of the variance ratios computed for 10 representative TFs in 10 representative tissues. The variance ratio is defined as the difference between the variance of the observed activity and that of the permuted activity, as a fraction of the former.

### Age-dependent inter-individual variation in HMGB3 activity

Our model is designed to analyze variation in TF activity across individuals for a given TF and tissue. To validate its biological usefulness, we tested whether the TF activity estimated from the GTEx data showed any biologically meaningful relationship with age. The most statistically significant association from this analysis was that HMGB2 activity in skeletal muscle significantly declines with aging (Pearson correlation r = –0.329, p-value = 2.4×10^−19^) (**Figure 3a**). Consistently, it has been previously reported that aging is associated with a loss of HMGB2 which contributes to the development of osteoarthritis (OA) (Taniguchi et al., 2009), one of the most common musculoskeletal disorders, which is strongly linked to aging (Loeser et al., 2016). Other tissues also showed age-dependent HMGB2 activity, including two adipose tissues (**Figure 3b**). Indeed, HMGB2 is known to play an important role in adipogenesis (Chen et al., 2021). Several other TFs also showed a significant dependence on age in these tissues (**Supplemental Figure S4a**). The comparison with age as a biological variable provided a natural opportunity to assess the added value of inferring TF activity using a model at the level of gene pairs as opposed to one at the level of individual genes (see **Methods**). Indeed, while the respective inferred HMGB2 activities are significantly correlated across skeletal muscle samples (**Supplemental Figure S4b**), the correlation with age is substantially less significant for HMGB2 activity as inferred using the individual-gene model (Pearson correlation r = –0.199, p-value = 1.02×10^−7^; **Supplemental Figure S4c**). This underscores the improved statistical power obtained with our gene-pair approach.

**Figure 3:**
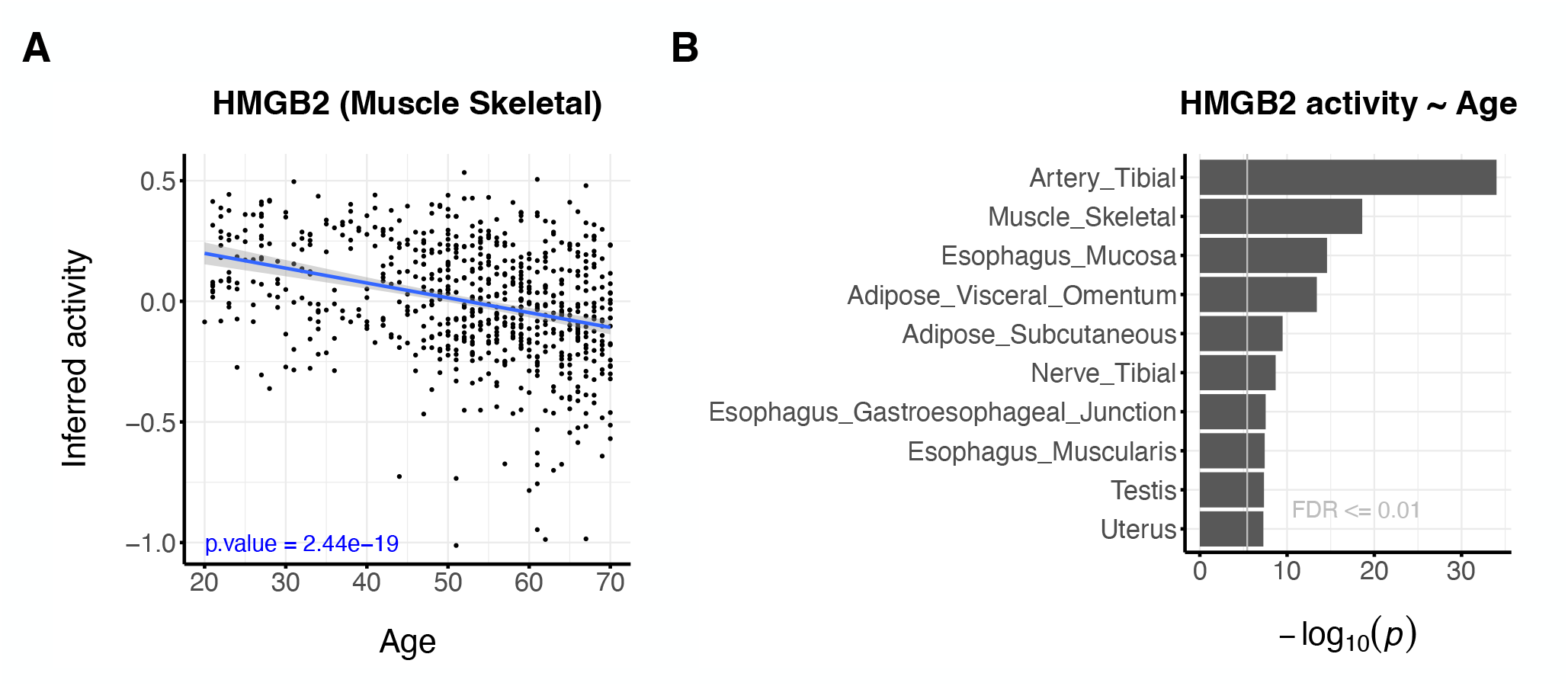
Inter-individual variation in TF activity. (a) Age-related HMGB2 activity across skeletal muscle samples in GTEx (p value = 2.44×10^−19;^; Pearson correlation test). (b) The ten tissues showing the strongest age-dependence of HMGB2 activity according to our method.

### Mapping genetic determinants of TF activity (aQTLs)

Our ability to estimate individual-specific TF activity, which can be viewed as a quantitative trait with a potential genetic component, provides an opportunity to use a standard genome-wide association study (GWAS) approach to map activity quantitative trait loci (aQTLs). These would be genetic modulators of the degree to which the TF proteins contributes to the expression level of its target genes in each individual in the GTEx cohort. To map aQTLs, we used a linear regression model in which major technical and biological covariates are included (see **Methods**). The most significant aQTL association in our analysis was for seen for rs146434626 and HMGN3 activity in frontal cortex (see **Supplemental Figure S5** for genome-wide profile), at a level of significance (p-value = 6.5×10^−16^) far exceeding the standard criterion for genome-wide significance (p-value < 5×10^−8^).

**Figure 4a** shows an overview of the aQTLs our method identified across all TFs and tissues, in which all variants with a p-value below the genome-wide significance threshold (p<5×10^−8^) have been marked. The standard significance threshold does not take into account the large number of TF/tissue combinations (total 55×49 = 2,695) that are being analyzed in parallel. However, many TFs have highly correlated inferred activities across tissues (**Supplemental Figure S6**). Thus, the effective number of independent traits tested is not merely the number of TF/tissue combinations, and care must be taken to not correct too harshly for multiple testing of traits and tissues in addition to genetic variants. We resorted to a permutation strategy to empirically estimate the false discovery rate (FDR) associated with to a particular p-value cutoff (see **Methods**). At the standard cutoff for a single trait (p<5×10^−8^), the FDR is rather high (∽70%), but it starts to drop as more stringent p-value thresholds are applied (**Figure 4b**). At the much more stringent cutoff of p<10^−11^, we discover 111 aQTLs (see **Supplemental Table S2** for details), at an FDR estimated to be around 25%. That we were able to discover such a high number of loci despite our a population size being in the hundreds, a couple of orders of magnitude smaller than the usual size in traditional GWAS, is a reflection of a high genetic component and a high signal-to-noise ratio reflected in our inferred TF activity trait. While we did not observe any tissue-sharing for these aQTLs, we did observe that a high correlation between the genetic effect on TF activity between similar tissues for variants that pass a lenient significance criterion (p<10^−6^) in one of tissues (**Supplemental Figure S7**). Thus, we would expect to observe more tissue sharing at larger population sizes with increased statistical power.

**Figure 4:**
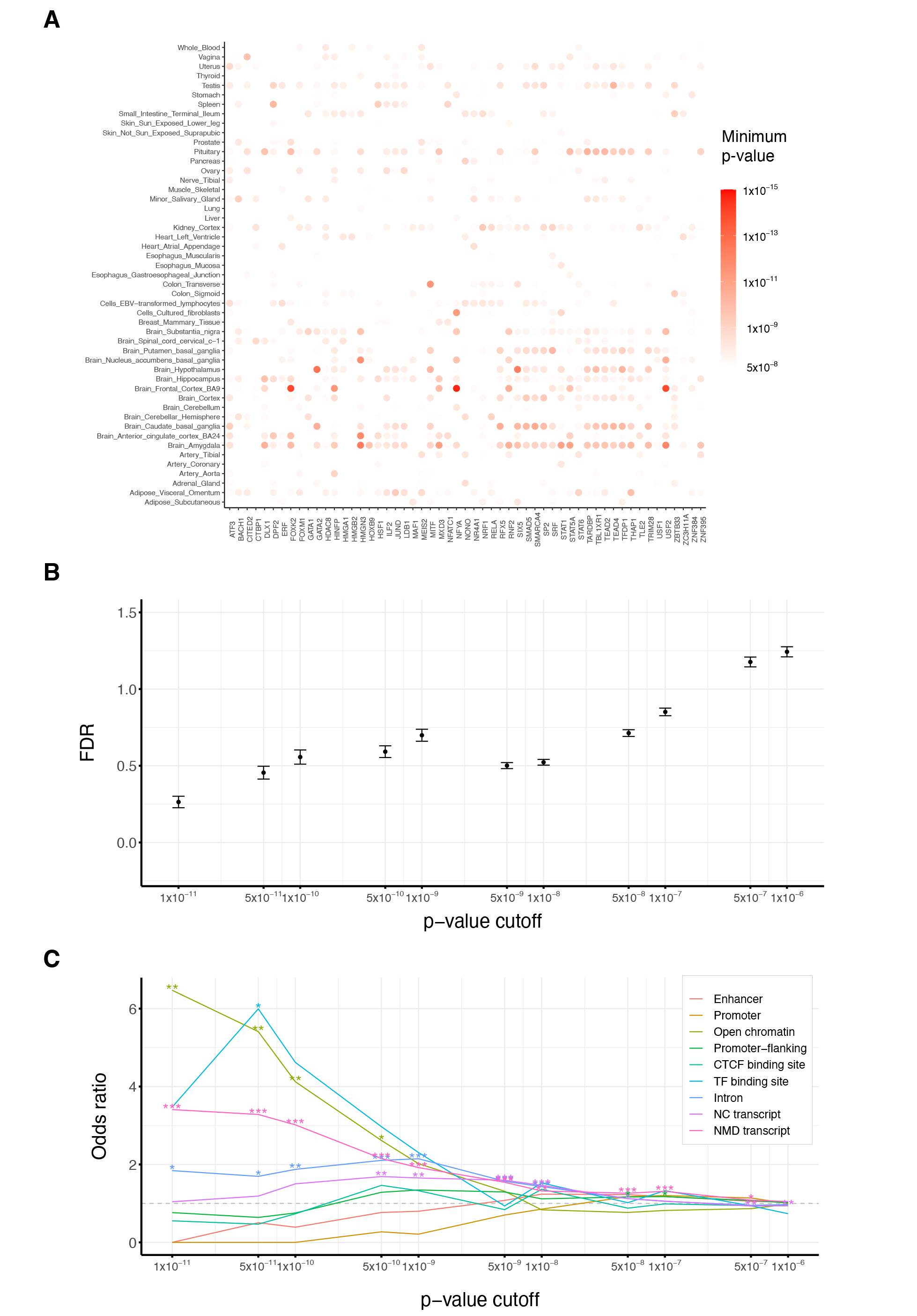
False discovery rate and functional enrichment for aQTLs discovered at different p-value cutoffs. (a) A heatmap showing the most significant p-value for each combination of trait TF and tissue. Only variants with p-values below 5×10^−8^ are indicated. (b) Plot showing the mean and SEM of FDRs estimated based on 100 random permutations. (c) Functional annotation enrichment of fine-mapped variants at various p-value cutoffs. Odds ratios and p-values were computed using Fisher’s exact test. Significance levels: *: p<=0.05, **: p<=0.01, ***: p<=0.001. NC: non coding, NMD: non mediated decay.

In addition to the loci significant at the threshold described above, we also explored hits at p<5×10^−8^ for putative corroborating biological mechanisms. One mechanism by which a protein encoded a gene at the aQTL locus might modulate the protein-level TF activity that is used as the trait is through direct protein-protein interaction as a transcriptional co-factor. An example of this is an aQTL on chromosome 18 (rs10775496 as the lead variant, p = 7.6×10^−10^), which our method identifies as a putative genetic determinant of the regulatory activity of RELA in cerebellar hemisphere tissue (**Figure 5a,b**). The nearest gene is SMAD4, and one of the most plausible causal variants (rs12456284; p = 7.8×10^−10^) resides in its 3’-UTR (**Figure 5c**). RELA, also known as p65, encodes a main subunit of NFkB, and it has been reported that RELA physically interacts with SMAD4 (Hirata-Tsuchiya et al., 2014).

**Figure 5:**
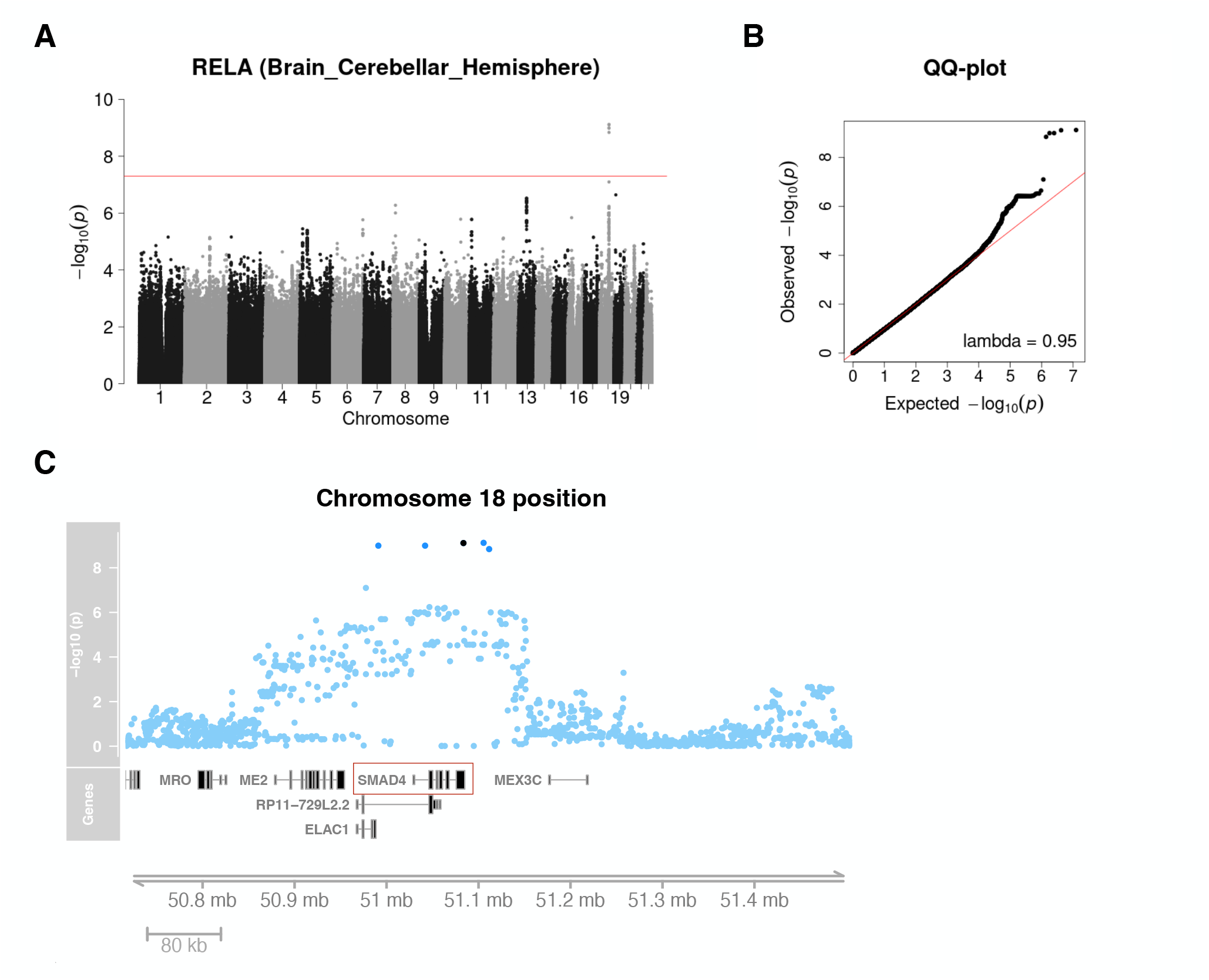
aQTL mapping results for transcription factor RELA in brain cerebellar hemisphere tissue. (a) Overview of genome-wide association analysis, using RELA activity as inferred by our GLM method as a quantitative trait. The red line indicates the p=5×10^−8^ significance level. (b) Quantile plot corresponding to panel (a), showing enrichment of small p-values. (c) Detailed view of one of the aQTL loci on chromosome 18. The dark blue points represent fine-mapped loci. The black point shows a 3’UTR variant of SMAD4, the transcript of which is highlighted in a red box.

To assess whether the identified aQTLs in aggregate have features indicative of gene regulation, we first fine-mapped aQTLs discovered at less stringent criteria (see **Methods**), and then at various p-value cutoffs computed enrichment ratios associated with different functional categories, according to a genome-wide functional annotation based on a widely used human cell line (see **Methods**). At more stringent p-value cutoffs, the identified variants are significantly enriched for open chromatin as well as TF binding sites (**Figure 4c**). This implies that, to a significant extent, the cascade of effects that ultimate leads to modulation of the target genes of the TF whose activity is analyzed, involves cis-regulatory changes at the aQTL locus, which presumably affect the expression level of a nearby “aGene” that mediates these effects.

As further validation of our aQTL mapping strategy, we asked whether there is a trend for the trait TF protein to have a known functional association with the protein product of the putative aGene taken as the gene closest to the aQTL. For each fine-mapped aQTL above the standard significance threshold, we determined which protein-coding gene had a transcription start site closest to the locus, and searched for known associations with the trait TF based on any type of evidence included in the STRING database (Szklarczyk et al., 2019). We found that such prior TF-aGene associations (**Supplemental Table S3**) are indeed more prevalent than expected by random chance (Fisher’s exact test, odds ratio = 1.83, p-value = 2.5×10^−6^). When the evidence is limited to coexpression, the association is less significant (odds ratio = 1.73; p-value = 0.002), whereas for protein interactions supported by experimental data it is slightly more significant (odds ratio 2.42; p-value = 5.0×10^−7^). Furthermore, we also observed that some aQTLs colocalize with a cis-eQTLs in the same tissue, which suggests the cis-target gene of the aQTLs may be the same as the “eGene” that was the trait for the eQTL association. For example, an aQTL (rs11547207) of HDAC8 in Adrenal Gland is colocalized with a cis-eQTL in the 5’-UTR of the CIB2 gene discovered in the GTEx project (see **Methods** and **Supplemental Figure S8**). Although in most cases the underlying mechanisms between mapped cis-target genes and the TFs are not obvious, such analysis can provide a starting point for the dissection of the mechanisms underlying trans-acting genetic effects.

## DISCUSSION

In this paper, we started by developing a robust general method for estimating protein-level TF activity levels from RNA-seq count data in a sample-specific manner. Our modeling is innovative in that we only focus on variation in the gene expression ratio between neighboring genes. This elegantly circumvents the confounding effect of sample-to-sample variation in chromatin structure on gene expression. We can attribute variation in genome-wide expression among GTEx samples to a particular TF by leveraging CRISPRi perturbation data. Using such an empirical signature as a predictor in our model has the advantage that context-dependent effects of functional versus non-functional TF binding, as well as indirect effects due to transcriptional cascades, are implicitly accounted for. On the downside, any TF perturbation signature is dependent on the cell line in which it was profiled, so our approach implicitly assumes that the effect of TF activity modulation is similar enough between that cell line and the GTEx samples that we analyze using our GLM.

In the future, using TF response signatures derived from large-scale multiplexed TF perturbation experiments coupled with single-cell expression profiling in other cell types (Schraivogel et al., 2020) (Replogle et al., 2021) might be used to refine our approach. While in our pioneering aQTL study in yeast we successfully used DNA binding specific models to simply predict TF responsiveness from upstream promoter sequence as a sliding-window sum of affinities (Lee & Bussemaker, 2010), this sequence-based strategy remains much more challenging in mammals, where cis-regulatory regions can be much farther away from a gene’s TSS, and where the combinatorial logic of TF binding is more complex.

Trans-acting variants that influence distal gene expression (i.e., trans-eQTLs) have been extensively studied and shown to be highly tissue-specific (Grundberg et al., 2012) (Brynedal et al., 2017) (Võsa et al., 2021). These variants can influence multiple genes by acting on regulatory circuits, and mapping them, and relating them explicitly to regulation by transcription factors, can help to clarify underlying mechanisms (Brynedal et al., 2017) (Võsa et al., 2021). With the TF activity inferred by our model and the parallel genotype data available, we were able to systematically identify genetic effects on TF activity in human tissues, regardless of whether the mRNA expression level of the gene encoding the TF has a genetic component or not.

A recent study inferred adipose-specific TF activities using a regulon-based approach and associated them with genetic variants at cardiometabolic trait GWAS loci, suggesting aQTLs can help reveal molecular mechanisms mediating GWAS signals (Hoskins et al., 2021). However, a systematic survey of aQTLs in other human tissues has not yet been conducted. Moreover, our gene-pair based GLM approach to inferring TF activity from transcriptome data is conceptually and technically very different from the regulon-based approach of (Hoskins et al., 2021).

Our model for inferring sample-specific TF activities has many potential applications in addition to identifying aQTLs. For instance, compared to normal human tissues, differential expression between cancer samples can be challenging to interpret due to the presence of genomic instabilities. Our gene-pair approach to inferred TF activity might be robust in the face of such instabilities, since a duplication in a region that includes both genes in a pair should not affect the ratio between their expression values. Another application would be to analyze individual-specific drug responsiveness by fitting the model on GTEx samples using an empirical drug-response signature as a predictor in the model. Genetic variants associated with differential drug responsiveness across individuals could thus be identified using our approach. This would provide an insight into genetic effects on individual-specific responsiveness to drugs and could have potential in the context of precision medicine.

## METHODS

### Collection of RNA-seq data from the GTEx project

To obtain the data to be used as the dependent variable in our model, we used gene counts from RNA-seq data from GTEx release v8, which encompasses over 15,000 samples across 54 tissues and over 800 individuals. We used data of 49 tissues (**Supplemental Table S1**) which have adequate sample sizes. For each tissue, RNA-seq counts were down-sampled to the minimum number of reads across all pertinent samples to make the library size equal across individuals.

### Construction of TF perturbation signatures

To construct the TF perturbation signatures used as the independent variable in our model, we obtained RNA-seq data reporting on the effect of CRISPRi knockdown of each of 66 TFs in K562 cell lines from the ENCODE data portal (Dunham et al., 2012). Starting from the alignment files for control and treatment samples in BAM format, we used featureCounts v2.0.0 (Liao et al., 2014) and the gene annotation of the genome assembly GRCh38.p12 from GENCODE release 29 (Frankish et al., 2021) to generate count matrices for each sample. Then we used R (version 4.0.2) package DESeq2 v1.28.1 (Love et al., 2014) to perform differential gene expression analysis for each CRISPRi RNA-seq experiment. After performing gene-level and sample-level quality control, we retained signatures for 55 TFs (**Supplemental Table S1**). The shrinkage log2 fold changes (Zhu et al., 2019) for each gene were used to define the perturbation signature for a given TF.

### Selection of gene pairs

To set up the design matrix for our model, we selected pairs of neighboring genes based on their location on the chromosome in GENCODE release 29. For each chromosome, we indexed all protein-coding genes according to their order along each chromosome, without regard for their transcription direction, and for each gene selected the next nearest gene as its paired gene. The resulting pairs were then filtered based on the distance between their respective annotated transcription start sites, which we required to be within the range of 10kb to 200 kb (**Supplemental Figure S2a**). For each tissue, we also left out pairs for which both genes had zero counts in all individuals. To assess the robustness of our model, we used an alternative way to generate two separate design matrices based on non-overlapping pairs of genes alone. Here, the first gene in a pair was required to have either an odd or an even index (“odd pairs” or “even pairs”). To analyze the ability of our nearest-neighbor design to account for variation in chromatin context along the genome, we also used sets of 5^th^ or 10^th^ nearest neighbor gene pairs to define the design matrix. In these cases, the TSS distance for each pair was typically greater than 200kb, and we did not impose a maximum-distance cutoff. As a further control, we also selected pairs in which each gene was randomly selected from different chromosome.

### Inference of TF activity based on gene-pair model

To estimate sample-specific (differential) TF activities, we used a generalized linear model based on the beta-binomial distribution:

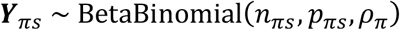

Here *π* = (*g*_1_, *g*_2_) denotes a gene pair. 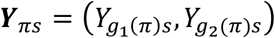 denotes the pair of RNA-seq counts *Y*_*gs*_ for gene pair *π* in sample *s*, with *g*_1_(*π*) and *g*_2_(*π*) defined as pair-to-gene mappings for the first and second gene in the pair, respectively. In the beta-binomial distribution, 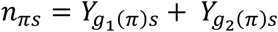 plays the role of the sample size, and *ρ*_*π*_ is the over-dispersion parameter. The binomial success rate *p*_*πs*_ was modeled as a function of the TF perturbation signature *χ*_*φg*_ (defined as the log_2_-ratio of the response of gene *g* each to perturbation of transcription factor *φ*) as follows:

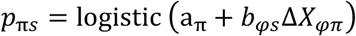

Here the intercept *a*_*π*_ models the mean expression ratio across samples between the two genes in each pair (which is not of interest by itself). 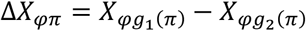 is the predictor based the TF perturbation signature for transcription factor *φ*, and *b*_*φs*_ the corresponding regression coefficient, which will be interpreted as an inferred, sample-specific, protein-level TF activity. A separate fit was performed for each TF for which a signature was available. The parameters *b*_*φs*_, *b*_*φs*_ and *ρ*_*π*_ were estimated by likelihood maximization, implemented in TensorFlow v2.9.1 (Abadi et al., 2016) and Python v3. For each tissue, a single joint fit to all individual samples for a given tissue was performed, allowing the over-dispersion parameter for each gene pair to be estimate without making any further assumptions. RNA-seq counts were down-sampled to make total counts the same as the smallest library size across all samples. Since the coefficients *b*_*φs*_ are defined up to an overall additive constant, we centered them at zero. To obtain a null distribution for the inferred TF activity *b*_*φg*_, the predictor *χ*_*φπ*_ was permuted randomly and independently among all gene pairs for each sample. The pair-level model was then fit to unpermuted expression counts ***Y***_*πg*_.

### Inference of TF activity based on individual-gene model

To model the count of individual genes across samples, we fit a generalized linear model with negative binomial distribution:

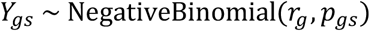

Here *Y*_*gs*_ denotes the RNA-seq count for gene *g* in sample *s*. The probability parameter *p*_*gs*_ is given by

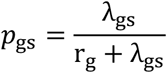

in terms of the gene-specific over-dispersion parameter *r*_*g*_ and the expected value *λ*_*gs*_ of Y_*gs*_, which in turns is parameterized as

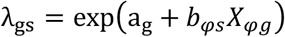

Here a_*g*_ is a gene-specific intercept that absorbs the variation in mean expression across samples, and *b*_*φs*_ is a sample-specific regression coefficient associated with the same perturbation-signature based predictor *χ*_*φg*_ as above (but now used at the individual-gene level), which we again interpret as an inferred sample-specific TF activity. The parameters *b*_*φs*_, *b*_*φs*_ and *r*_*g*_ were estimated by likelihood maximization as above. For computational efficiency, we optionally set a_*g*_ to the actual mean

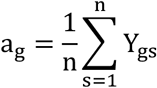

rather than treating it as a fit parameter.

### Mapping aQTLs using genome-wide association analysis of inferred TF activity

Treating the inferred activity level *b*_*φs*_ for a given transcription factor *φ* as a sample-specific quantitative trait, we computed their association with genetic variants using a standard genome-wide association (GWA) analysis based on univariate linear regression using the fastGWA tool from GCTA v1.93.1 (Yang et al., 2011). To account for non-genetic sources of variation, we included as covariates unobserved confounding variables also used in the standard eQTL analysis in the GTEx project (THE GTEX CONSORTIUM, 2020), including five genotype-based principal components that account for population structure; the genotyping platform; the biospecimen source site; the cohort to which the sample belonged (organ donor or postmortem); sex; and age. The association study of TF activity traits was done for all TF and tissue combinations.

### Statistical fine-mapping

We uniformly processed the summary statistics of association tests for each TF activity in a given tissue for statistical fine-mapping. We first used GCTA-COJO v1.93.1 (Yang et al., 2011) (Yang et al., 2012) to identify conditionally independent lead variants (pJ < 5 × 10^−6^) for each trait. We then analyzed all variants within the ±1Mb region of independent variants with FINEMAP v1.3.1, a Bayesian fine-mapping method (Benner et al., 2016). The covariance matrices required by FINEMAP were generated by PLINK v2.0 (Purcell et al., 2007) and LDstore v1.1 (Benner et al., 2017). The variants with a high Bayes factor (log_10_(BF) ≥ 2) were then retained for a set of plausibly causal variants.

### Empirical false discovery rates

We adopted a permutation strategy to estimate the false discovery rate (FDRs) at a given p-value cutoff. In each permutation, the association between the genotype and the TF activity traits across individuals was randomized, while preserving both the correlation structure among different TFs and that among genetic variants, as well as the covariates. We repeated the GWA analysis after each of 100 random permutations, for each tissue separately. For any given p-value cutoff, this allowed us to estimate the mean expected number of false discoveries across all combinations of genetics variant, TFs, and tissue. The FDR was then computed as the ratio of this expected mean and the actual number of discoveries at the same p-value cutoff.

### Enrichment of aQTLs for specific genomic annotations

We first performed functional annotation of all fine-mapped genetic variants as described above using the ENSEMBL Variant Effect Predictor (VEP) tool (McLaren et al., 2016). To test for functional enrichment among the set of fine-mapped aQTLs discovered across TFs and tissues at a given p-value cutoff, we used Fisher’s exact test.

### Enrichment of protein association annotations between the TF and the nearest genes to the aQTL

We downloaded functional protein associations from the STRING v11.5 database (Szklarczyk et al., 2019), where evidence types encompass conserved neighborhood, co-occurrence, fusion, coexpression, experiments, curated databases, and text mining. For each fine-mapped aQTL above the standard significance threshold (5×10^−8^), we posited that the protein-coding gene with a transcription start site closest to the locus was the “aGene”. We collected all unique TF-aGene pairs, and then used Fisher’s exact text to determine whether these TF-aGene pairs are enriched in terms of prior annotated protein associations, either in aggregate or in each separate category.

### Colocalization of aQTLs and cis-eQTLs

We obtained the fine-mapped GTEx cis-eQTLs and summary statistics from the eQTL Catalogue (Kerimov et al., 2021). For each candidate variant in the intersection of the set of fine-mapped aQTLs and the set of eQTLs mapped in the same tissue, we performed colocalization analysis with the R package “coloc” (Giambartolomei et al., 2014) using a 1Mb window around the corresponding variant, and computed a posterior probability of colocalization (PP4). We then used the R package “LocusCompareR” (Liu et al., 2019) to visualize of aQTL-eQTL colocalization events.

### Code and Data Availability

All our scripts are available at github.com/xl27/GTEx_aQTLs. Summary statistics and fine-mapping results of all aQTL analyses are available at bussemakerlab.org/papers/Li-aQTL

## ACKNOWLEDGEMENTS

This research was supported by NIH award R01MH106842 (to H.J.B. and T.L.) and a PhRMA foundation pre-doctoral fellowship in informatics to X.L. We thank Dr. Chaitanya Rastogi and Dr. H. Tomas Rube for critical advice on algorithm development, Dr. Júlia Domingo and Dr. John Morris for valuable suggestions on analysis of genetic variants, and Dr. Pejman Mohammadi as well as members of Bussemaker and Lappalainen labs for helpful discussions.

## AUTHOR CONTRIBUTIONS

H.J.B and X.L. developed the methodology; X.L. wrote all the software and performed all analyses, under the supervision of H.J.B and T.L; X.L and H.J.B wrote the manuscript; all authors edited and approved the manuscript.

## DECLARATION OF INTERESTS

The authors declare no competing interests.

## SUPPLEMENTAL INFORMATION

**Supplemental Table S1: Transcription factors and tissues analyzed in this study**.

**Supplemental Table S2: Summary statistics for aQTLs discovered at a p-value threshold of 10**^**-11**^. **Supplemental Table S3: aQTLs annotated by prior protein-protein association with nearest gene**.

**Supplementary Figure S1**: **Overall framework of the TF activity inference method**. (a) We performed the differential analysis of CRISPRi RNA-seq experiments targeting a single TF and used the shrinkage-based log_2_-fold-change to define the genome-wide TF perturbation response signature. (b) Schematic diagram showing how the predictor was permuted across gene pairs, for each sample independently. The same model fit was then used to construct a null distribution for the inferred TF activities.

**Supplemental Figure S2: Analyzing the robustness of our TF activity estimation method**. (a) Cutoffs on distances between TSS for neighboring pairs used, shown relative to their distribution. (b) Construction of odd pair set and even pair set (c) An example of consistency between activities inferred from odd pair set and even pair set. (d) Distribution of correlations between odd pair set and even pair set for 10 representative TFs in 10 representative tissues.

**Supplemental Figure S3: Comparison of gene-level model and pair-level model with different sets of pairs**. (a) Construction of sets of pairs with varying distances. (b) Ecdf plot of distances for different pair sets. (c) Comparison of activities inferred using the gene-level model or the pair-level models with different sets of pairs. We investigated if modeling on the neighboring pairs can capture more local effects, by developing a “gene-level” model (Methods) and “pair-level” models with the distances between genes within a pair varied (Methods), and comparing their inference results with the neighboring pair-level model. In the gene-level model, we fit a GLM with negative binomial distribution on the count data of individual genes across samples for each tissue. To further investigate if the distance for genes within a pair would have an impact on the inference, we selected pairs that are further away from each other, in addition to neighboring gene pairs. For instance, we constructed 5^th^ gene pairs, 10^th^ gene pairs, and random gene pairs selected from different chromosomes. Comparing the inferred activity, among pair-level models with different sets of gene pairs, the gene-level model is the most correlated with the random pair set and the least correlated with the neighboring pair set. Besides, we found neighboring pair set has the highest correlation to the 5th pair set and the lowest correlation to the random pairs (Suppl Fig 2c). The fact that the neighboring pair-level model shows the most different results is consistent with our expectation since it captures more local effects.

**Supplemental Figure S4: Age dependence of TF activity across all tissues**. (a) A heatmap showing the statistical significance of the Pearson correlation between age and TF activity inferred from the pair-level model, computed for all TFs across all tissues. (b) Correlation between gene-pair and individual-gene model based estimates of HMGB2 activity across skeletal muscle samples. (c) Age dependence of HMGB2 activity in skeletal muscle as inferred using individual-gene model.

**Supplemental Figure S5: aQTL mapping results for HMGN3 in brain frontal cortex tissue**. The most significant aQTL association in our analysis

**Supplemental Figure S6: Correlation in inferred activity between TFs across individuals**. The heatmap shows the Pearson correlation between profiles of inferred activity in skeletal muscle for each TF pair.

**Supplemental Figure S7: Tissue sharing of aQTL effect sizes**. (a) An example showing the correlation in effect size (regression coefficient in linear model in which the genotype of the independent variable) between two adipose tissues for all TF-aQTL pairs discovered at p-value < 1×10^−6^. (b) Heatmap showing how tissues cluster based on the correlations shown in panel (a).

**Supplemental Figure S8: Colocalization of an aQTL and cis-eQTL in Adrenal Gland**. An example of colocalization between aQTL and eQTL for the variant rs11547207 in Adrenal Gland. The right panel shows the 1Mb region surrounding the lead variant, using either aQTL or eQTL summary statistics. The left panel shows the correlation between the respective p-values for all variants within the same 1Mb region. PP4: posterior probability of colocalization from COLOC.

## Notes

### Competing Interest Statement

The authors have declared no competing interest.

