## Supplemental Figures for "Identifying Genetic Regulatory Variants that Affect Transcription Factor Activity"

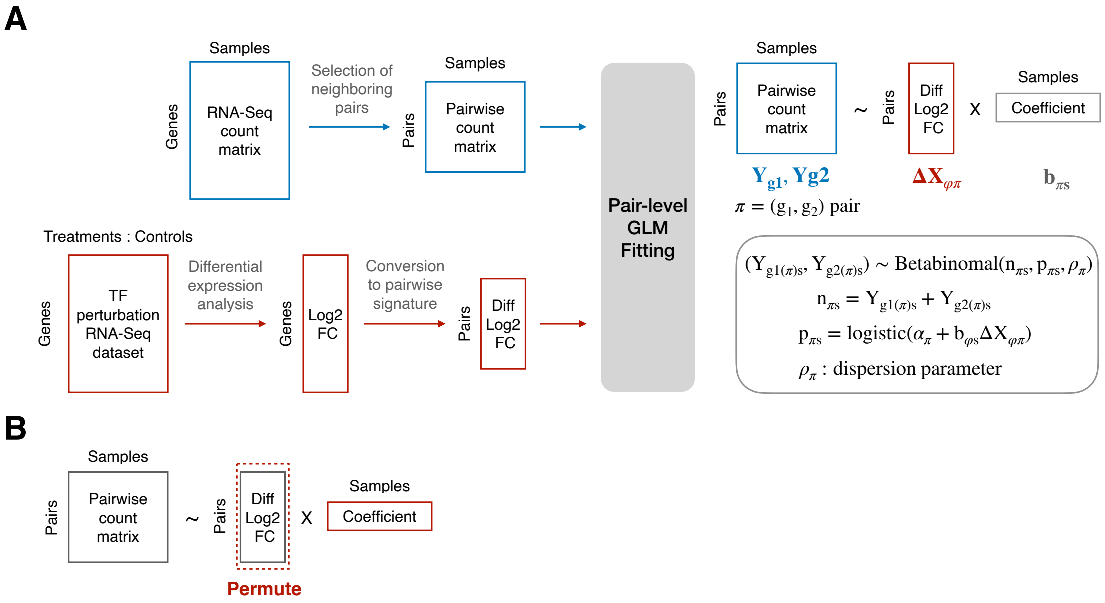


**Figure S1**: **Overall framework of the TF activity inference method**. (a) We performed the differential analysis of CRISPRi RNA-seq experiments targeting a single TF and used the shrinkage-based $log_2$-fold-change to define the genome-wide TF perturbation response signature. (b) Schematic diagram showing how the predictor was permuted across gene pairs, for each sample independently. The same model fit was then used to construct a null distribution for the inferred TF activities.

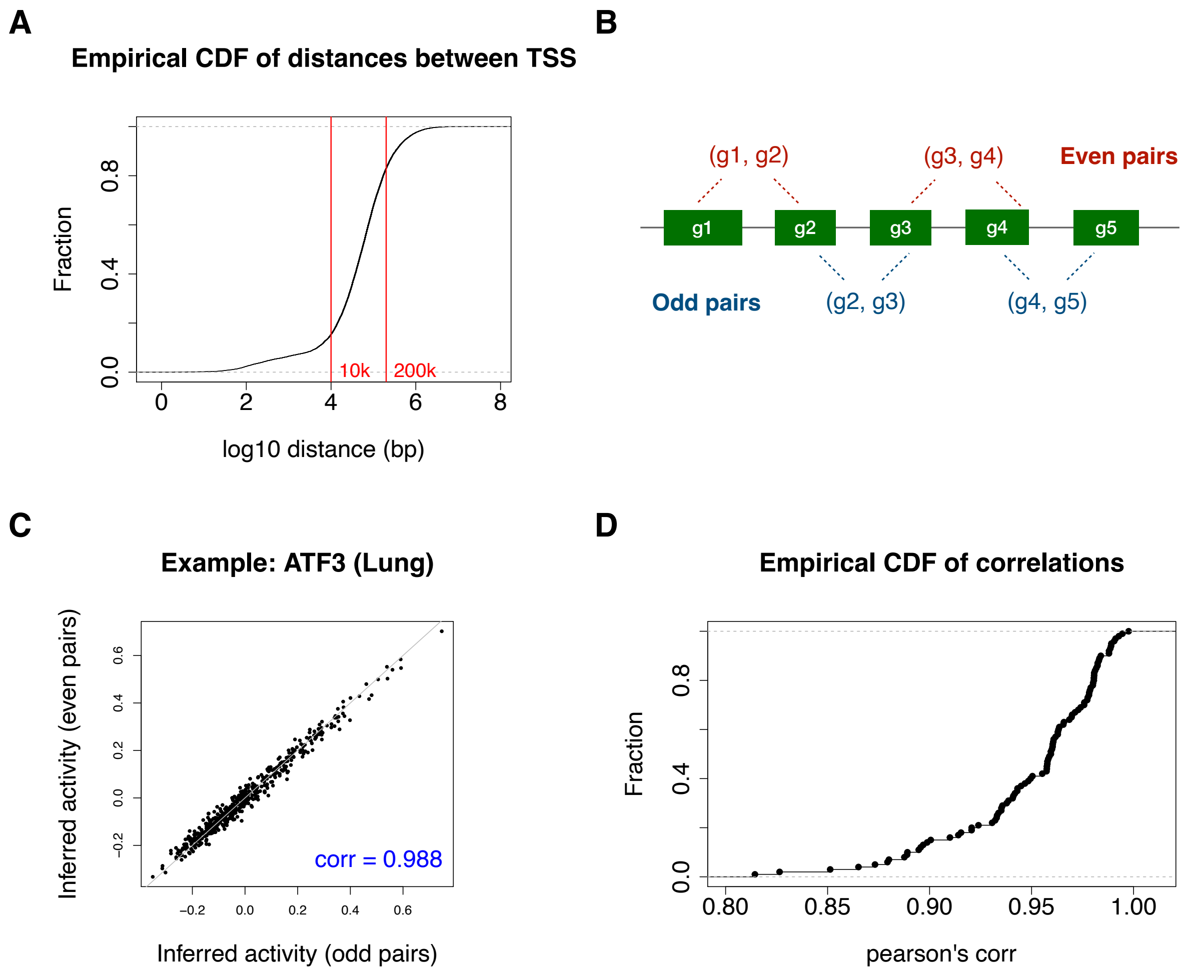


**Figure S2: Analyzing the robustness of our TF activity estimation method.** (a) Cutoffs on distances between TSS for neighboring pairs used, shown relative to their distribution. (b) Construction of odd pair set and even pair set (c) An example of consistency between activities inferred from odd pair set and even pair set. (d) Distribution of correlations between odd pair set and even pair set for 10 representative TFs in 10 representative tissues.

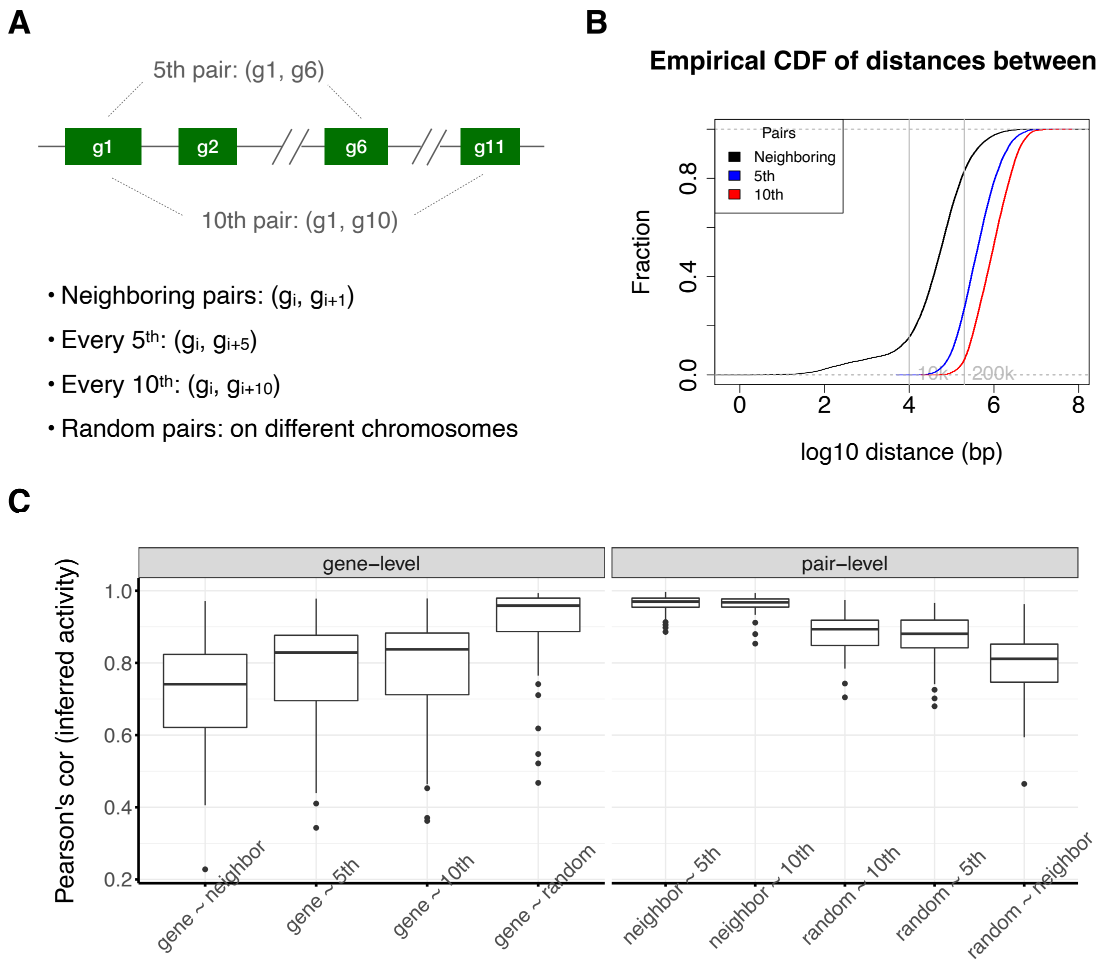


**Figure S3: Comparison of gene-level model and pair-level model with different sets of pairs**. (a) Construction of sets of pairs with varying distances. (b) Ecdf plot of distances for different pair sets. (c) Comparison of activities inferred using the gene-level model or the pair-level models with different sets of pairs. We investigated if modeling on the neighboring pairs can capture more local effects, by developing a "gene-level" model (Methods) and "pair-level" models with the distances between genes within a pair varied (Methods), and comparing their inference results with the neighboring pair-level model. In the gene-level model, we fit a GLM with negative binomial distribution on the count data of individual genes across samples for each tissue. To further investigate if the distance for genes within a pair would have an impact on the inference, we selected pairs that are further away from each other, in addition to neighboring gene pairs. For instance, we constructed $5^{th}$ gene pairs, $10^{th}$ gene pairs, and random gene pairs selected from different chromosomes. Comparing the inferred activity, among pair-level models with different sets of gene pairs, the gene-level model is the most correlated with the random pair set and the least correlated with the neighboring pair set. Besides, we found neighboring pair set has the highest correlation to the 5th pair set and the lowest correlation to the random pairs (Suppl Fig 2c). The fact that the neighboring pair-level model shows the most different results is consistent with our expectation since it captures more local effects.

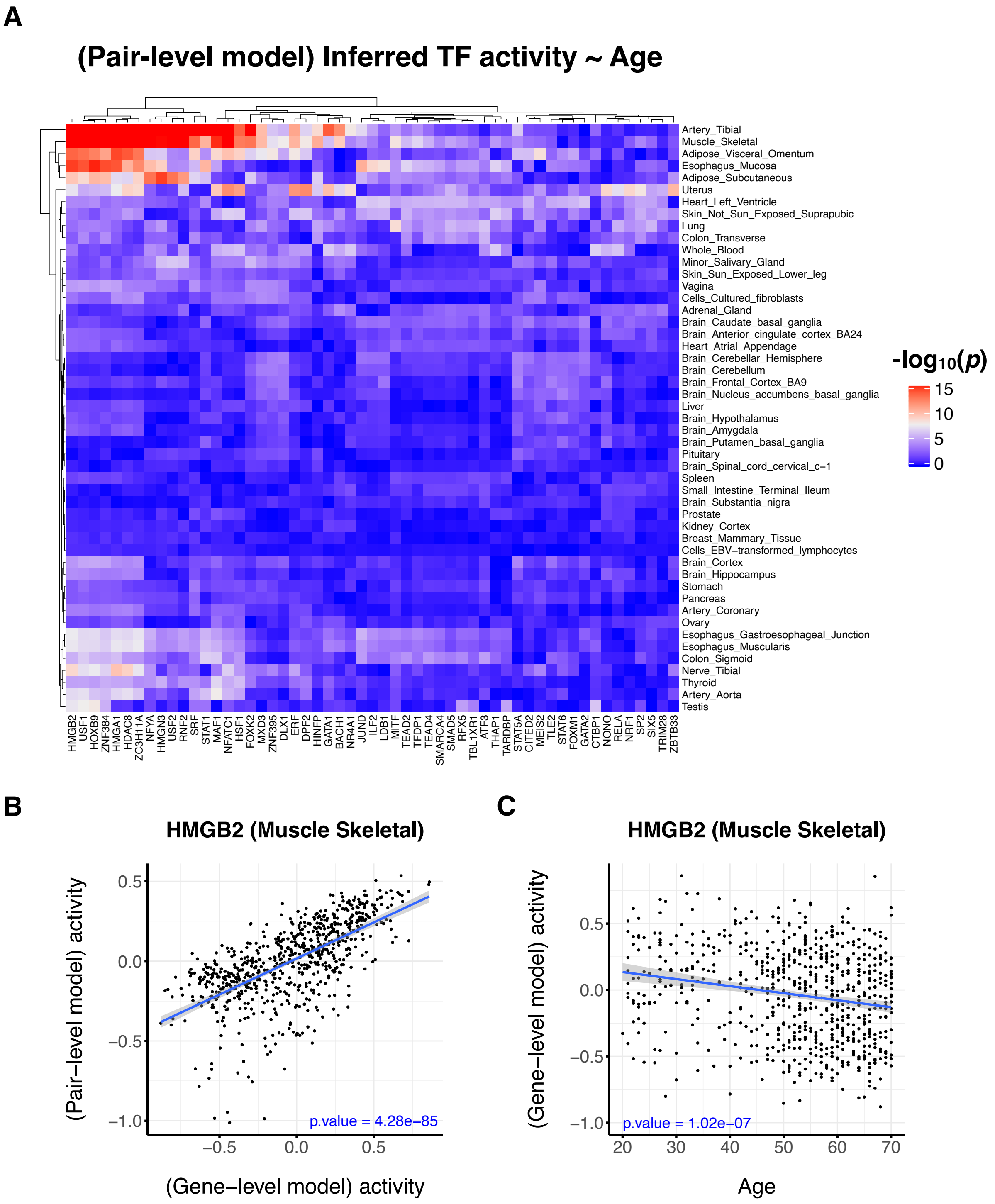


**Figure S4: Age dependence of TF activity across all tissues.** (a) A heatmap showing the statistical significance of the Pearson correlation between age and TF activity inferred from the pair-level model, computed for all TFs across all tissues. (b) Correlation between gene-pair and individual-gene model based estimates of HMGB2 activity across skeletal muscle samples. (c) Age dependence of HMGB2 activity in skeletal muscle as inferred using individual-gene model.

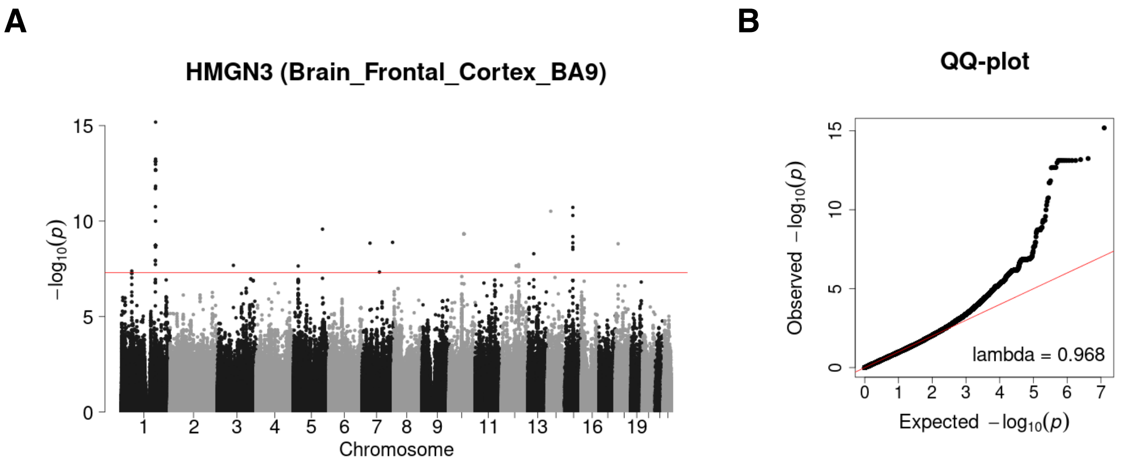


**Figure S5: aQTL mapping results for HMGN3 in brain frontal cortex tissue.** The most significant aQTL association in our analysis

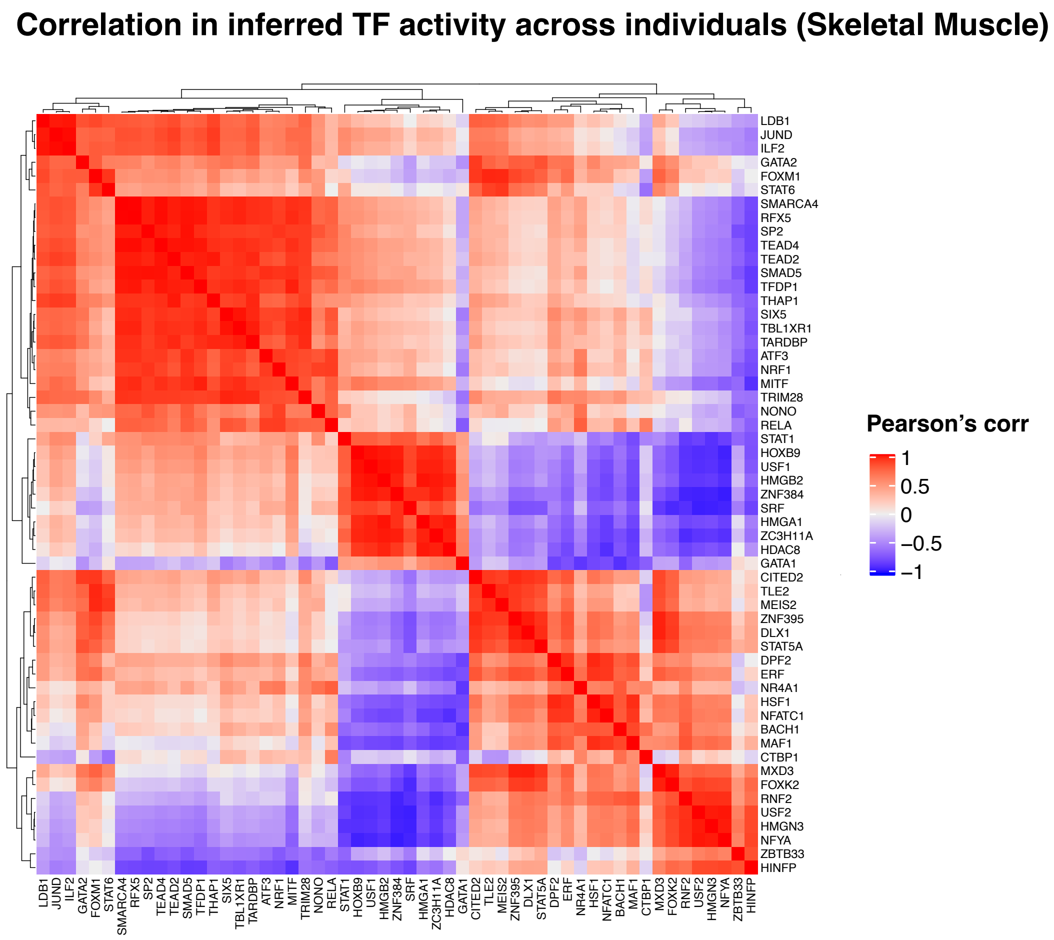


**Figure S6: Correlation in inferred activity between TFs across individuals.** The heatmap shows the Pearson correlation between profiles of inferred activity in skeletal muscle for each TF pair.

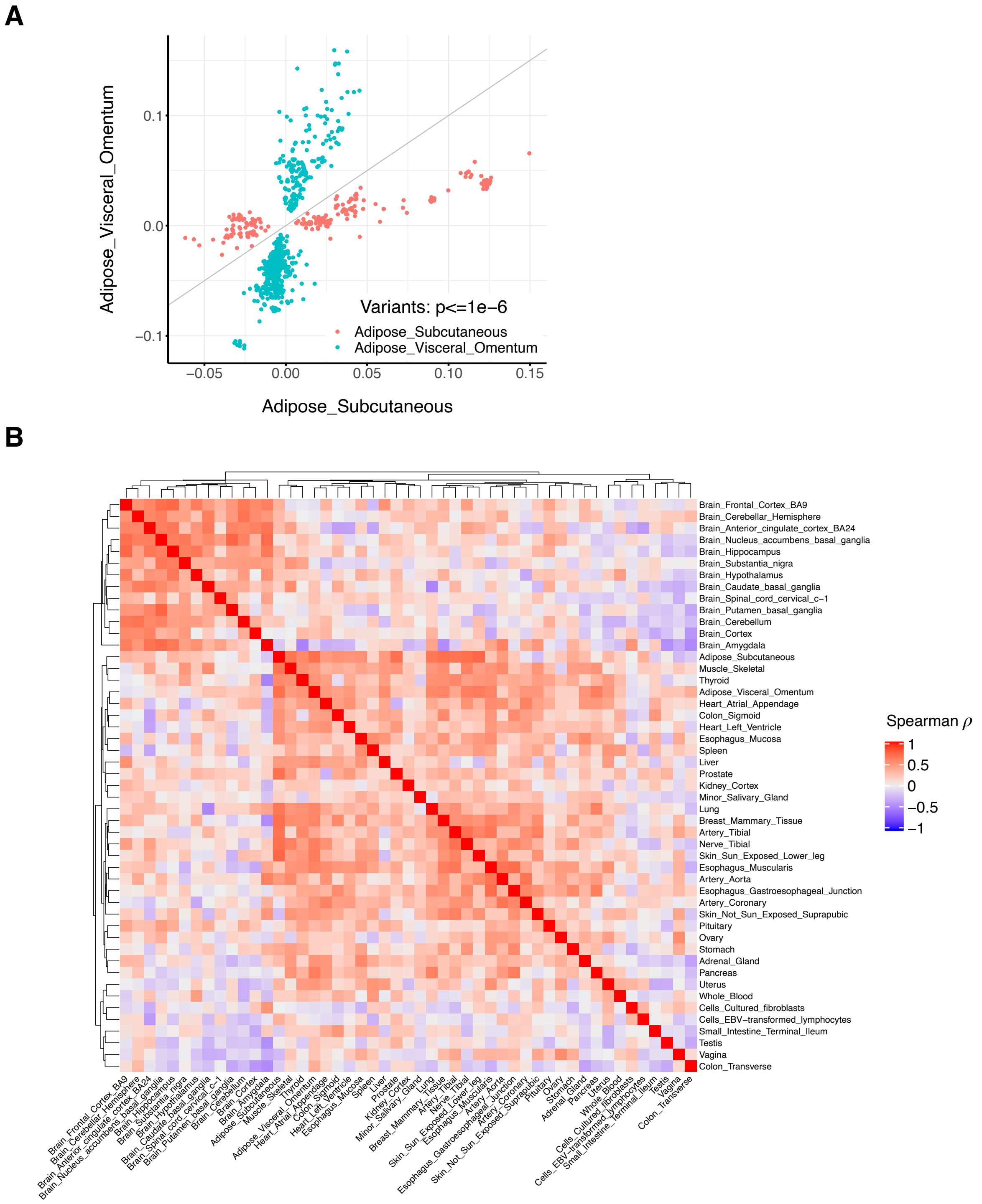


**Figure S7: Tissue sharing of aQTL effect sizes.** (a) An example showing the correlation in effect size (regression coefficient in linear model in which the genotype of the independent variable) between two adipose tissues for all TF-aQTL pairs discovered at p-value < $1x10^{-6}$. (b) Heatmap showing how tissues cluster based on the correlations shown in panel (a).

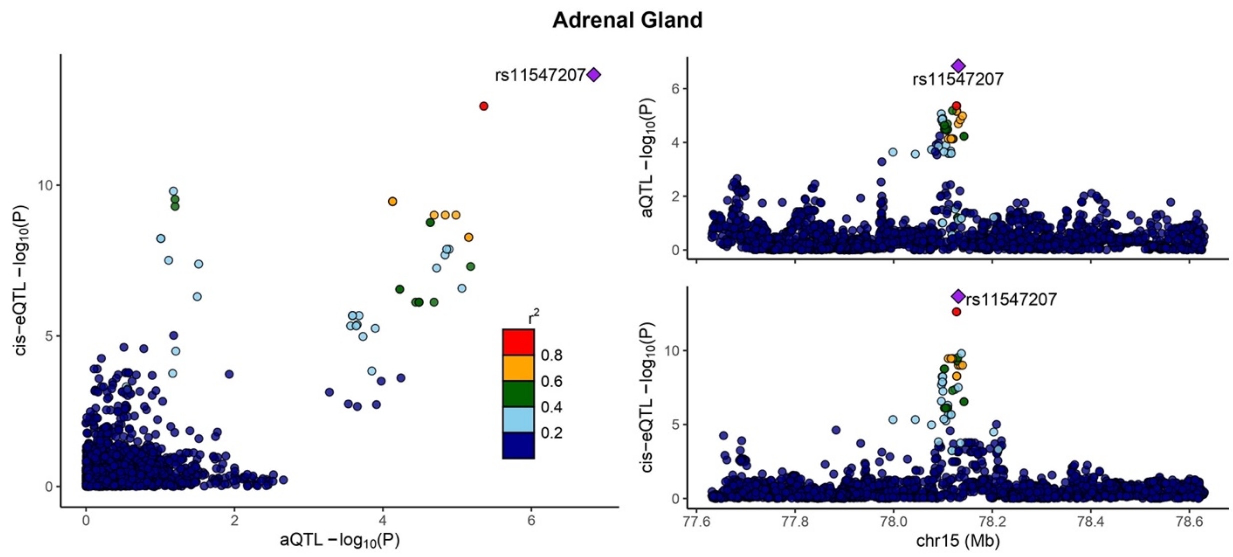


**Figure S8: Colocalization of an aQTL and cis-eQTL in Adrenal Gland.** An example of colocalization between aQTL and eQTL for the variant rs11547207 in Adrenal Gland. The right panel shows the 1Mb region surrounding the lead variant, using either aQTL or eQTL summary statistics. The left panel shows the correlation between the respective p-values for all variants within the same 1Mb region. PP4: posterior probability of colocalization from COLOC.

**Table S1: Transcription factors and tissues analyzed in this study.**

**TFs**

| | | | | |
|---|---|---|---|---|
| DLX1 | HOXB9 | USF1 | TFDP1 | RNF2 |
| NONO | NFATC1 | DPF2 | TARDBP | SMAD5 |
| TLE2 | HMGB2 | NR4A1 | HMGA1 | ZNF395 |
| NRF1 | STAT1 | NFYA | CTBP1 | GATA1 |
| ATF3 | SRF | ZBTB33 | ZC3H11A | ERF |
| TRIM28 | BACH1 | STAT5A | TBL1XR1 | FOXM1 |
| ZNF384 | MEIS2 | SIX5 | MAF1 | TEAD2 |
| FOXK2 | GATA2 | MITF | ILF2 | THAP1 |
| STAT6 | JUND | HSF1 | RFX5 | RELA |
| SMARCA4 | USF2 | MXD3 | HDAC8 | HINFP |
| HMGN3 | CITED2 | LDB1 | SP2 | TEAD4 |

**Tissues**

| | |
|---|---|
| Adipose_Subcutaneous | Esophagus_Mucosa |
| Adipose_Visceral_Omentum | Esophagus_Muscularis |
| Adrenal_Gland | Heart_Atrial_Appendage |
| Artery_Aorta | Heart_Left_Ventricle |
| Artery_Coronary | Kidney_Cortex |
| Artery_Tibial | Liver |
| Brain_Amygdala | Lung |
| Brain_Anterior_cingulate_cortex_BA24 | Minor_Salivary_Gland |
| Brain_Caudate_basal_ganglia | Muscle_Skeletal |
| Brain_Cerebellar_Hemisphere | Nerve_Tibial |
| Brain_Cerebellum | Ovary |
| Brain_Cortex | Pancreas |
| Brain_Frontal_Cortex_BA9 | Pituitary |
| Brain_Hippocampus | Prostate |
| Brain_Hypothalamus | Skin_Not_Sun_Exposed_Suprapubic |
| Brain_Nucleus_accumbens_basal_ganglia | Skin_Sun_Exposed_Lower_leg |
| Brain_Putamen_basal_ganglia | Small_Intestine_Terminal_Ileum |
| Brain_Spinal_cord_cervical_c-1 | Spleen |
| Brain_Substantia_nigra | Stomach |
| Breast_Mammary_Tissue | Testis |
| Cells_Cultured_fibroblasts | Thyroid |
| Cells_EBV-transformed_lymphocytes | Uterus |
| Colon_Sigmoid | Vagina |
| Colon_Transverse | Whole_Blood |
| Esophagus_Gastroesophageal_Junction | |
